# SmartPeak automates targeted and quantitative metabolomics data processing

**DOI:** 10.1101/2020.07.14.202002

**Authors:** Svetlana Kutuzova, Pasquale Colaianni, Hannes Röst, Timo Sachsenberg, Oliver Alka, Oliver Kohlbacher, Bo Burla, Federico Torta, Lars Schrübbers, Mette Kristensen, Lars Nielsen, Markus J. Herrgård, Douglas McCloskey

## Abstract

SmartPeak is an application that encapsulates advanced algorithms to enable fast, accurate, and automated processing of CE-, GC- and LC-MS(/MS) data, and HPLC data for targeted and semi-targeted metabolomics, lipidomics, and fluxomics experiments.

**Highlights:**

- Novel algorithms for retention time alignment, calibration curve fitting, and peak integration
- Enables reproducibility by reducing operator bias and ensuring high QC/QA
- Automated pipeline for high throughput targeted and/or quantitative metabolomics, lipidomics, and fluxomics data processing from multiple analytical instruments
- Manually curated data set of LC-MS/MS, GC-MS, and HPLC integrated peaks for further algorithm development and benchmarking

---

Analytical labs in pharmaceutical, biotech, forensic, clinical, and diagnostic sectors across the world are routinely required to provide highly accurate, reproducible, and quantitative amounts of small molecules and peptides in complex sample matrices for screening and quality control applications ^1,2^. In addition, academic and other research-based life science laboratories also require accurate, reproducible, and quantitative amounts of small molecules and peptides for quantitative modeling and analysis of physiology ^3^, and drug and biomarker discovery ^4^ applications. The gold standard instrumentations of choice for accurate, reproducible, and quantitative analysis include liquid chromatography (LC) or gas chromatography (GC) coupled to a single (Q) or triple quadrupole (QqQ) mass spectrometer (MS) operated in single ion mode (SIM) or single reaction monitoring (SRM) mode, respectively due to their combination of selectivity, sensitivity, linearity, and economy. However, as high resolution (HRMS) instrumentation technology improves, it is expected that quantitative workflows using sequential window acquisition of all theoretical fragment ion spectra (SWATH) and data independent acquisition (DIA) acquisition types will also become routine on these platforms (e.g., ^5,6^ for evaluations of quantitative performance using HRMS instruments compared to QqQ). In addition, high pressure liquid chromatography (HPLC) coupled to a variety of detection sources (e.g., UV, refractive index, etc.) are also commonly employed.

Improvements in instrumentation technology to perform targeted and non-targeted acquisition with quantitative or non-quantitative analysis of small molecules and proteins have increased the speed of running samples and the volume of data generated by each sample. Consequently, the bottleneck of many analytical labs has shifted from high throughput method development to high throughput data processing. Data processing is often done by commercial or vendor-specific software. However, most commercial and vendor-specific software were not written with high throughput data processing needs in mind. These needs include algorithms that are robust yet flexible to deal with heterogeneous and complex data, a graphical user interface (GUI) that supports configuring, monitoring, and reviewing big data workflows, and speed/performance when working with large data sets. Lack of these prerequisites often leads to long, tedious, and error prone manual data processing sessions. Free tools have emerged (e.g., Skyline^7^, MRMAnalyzer ^8^, MRMPROBS ^9^, MAVEN^10,11^, MaxQuant^12^, and XCMS-MRM ^13^) that address some of these needs, but often lack the features or robustness provided by commercial or vendor-specific software to allow for reproducible use in regulated environments.

To address the lack of robust algorithms and software suitable for scaling quantitative analytical data processing to Big Data regimes, we have developed a suite of algorithms that are a part of the OpenMS ^14^ toolkit (Table 1). In addition, we have developed an application for routine deployment of those algorithms called SmartPeak. SmartPeak provides graphical- and/or command-line-based user input validation, workflow configuration, data visualization and review, logging, and reporting. SmartPeak can be run on multiple operating systems or on cloud infrastructures. The workflow automates all steps from peak detection and integration, to calibration curve optimization, to quality control reporting (Fig 1). The workflow supports SIM, SRM, PRM, SWATH, and DIA data types derived from CE-, LC-, GC-MS(/MS) and HPLC instrumentation for applications in metabolomics, lipidomics, fluxomics, and proteomics.

**Table. 1.** An overview of classes, methods, and features added to the OpenMS library for targeted and quantitative metabolomics data analysis.

| Header file | Category | Contribution | Description |
| --- | --- | --- | --- |
| OpenMS/FORMAT/ChromeleonFile.h | IO | New class | ChromeleonFile |
| OpenMS/KERNEL/SpectrumHelper.h | Chromatogram manipulation | New methods | subtractMinimumIntensity and removePeaks |
| OpenMS/FORMAT/MRMFeatureQCFile.h | IO | New class | MRMFeatureQCFile |
| OpenMS/FORMAT/AbsoluteQuantitationMethodFile.h | IO | New class | AbsoluteQuantitationMethodFile |
| OpenMS/FORMAT/AbsoluteQuantitationStandardsFile.h | IO | New class | AbsoluteQuantitationStandardsFile |
| OpenMS/ANALYSIS/QUANTITATION/AbsoluteQuantitationMethod.h | Format | New class | AbsoluteQuantitationMethod |
| OpenMS/METADATA/AbsoluteQuantitationStandards.h | Format | New class | AbsoluteQuantitationStandards |
| OpenMS/ANALYSIS/OPENSWATH/MRMFeatureQC.h | Format | New structs | ComponentQCs, ComponentGroupQCs, ComponentGroupPairsQCs |
| OpenMS/ANALYSIS/MAPMATCHING/TransformationModel.h | Math | New feature | Parameter weighting |
| OpenMS/ANALYSIS/MAPMATCHING/TransformationModelLinear.h | Math | New feature | Parameter weighting |
| OpenMS/ANALYSIS/OPENSWATH/MRMFeatureSelector.h | Feature selection | New class | MRMFeatureSelector |
| OpenMS/ANALYSIS/OPENSWATH/MRMBatchFeatureSelector.h | Feature selection | New class | MRMBatchFeatureSelector |
| OpenMS/ANALYSIS/OPENSWATH/PeakIntegrator.h | Feature integration | New class | PeakIntegrator |
| OpenMS/ANALYSIS/OPENSWATH/MRMFeatureFilter.h | Feature filtering | New class | MRMFeatureFilter |
| OpenMS/ANALYSIS/OPENSWATH/MRMTransitionGroupPicker.h | Feature finding | New features | Peak integration and background identification/subtraction methods, metric calculations, and transition retention time and peak area reporting |
| OpenMS/ANALYSIS/QUANTITATION/AbsoluteQuantitation.h | Feature quantification | New class | AbsoluteQuantitation |
| OpenMS/MATH/MISC/EmgGradientDescent.h | Feature fitting | New class | EmgGradientDescent |
| SmartPeak/algorithm/MRMFeatureValidator.h | Feature validation | New class | MRMFeatureValidator |

**Fig. 1.**
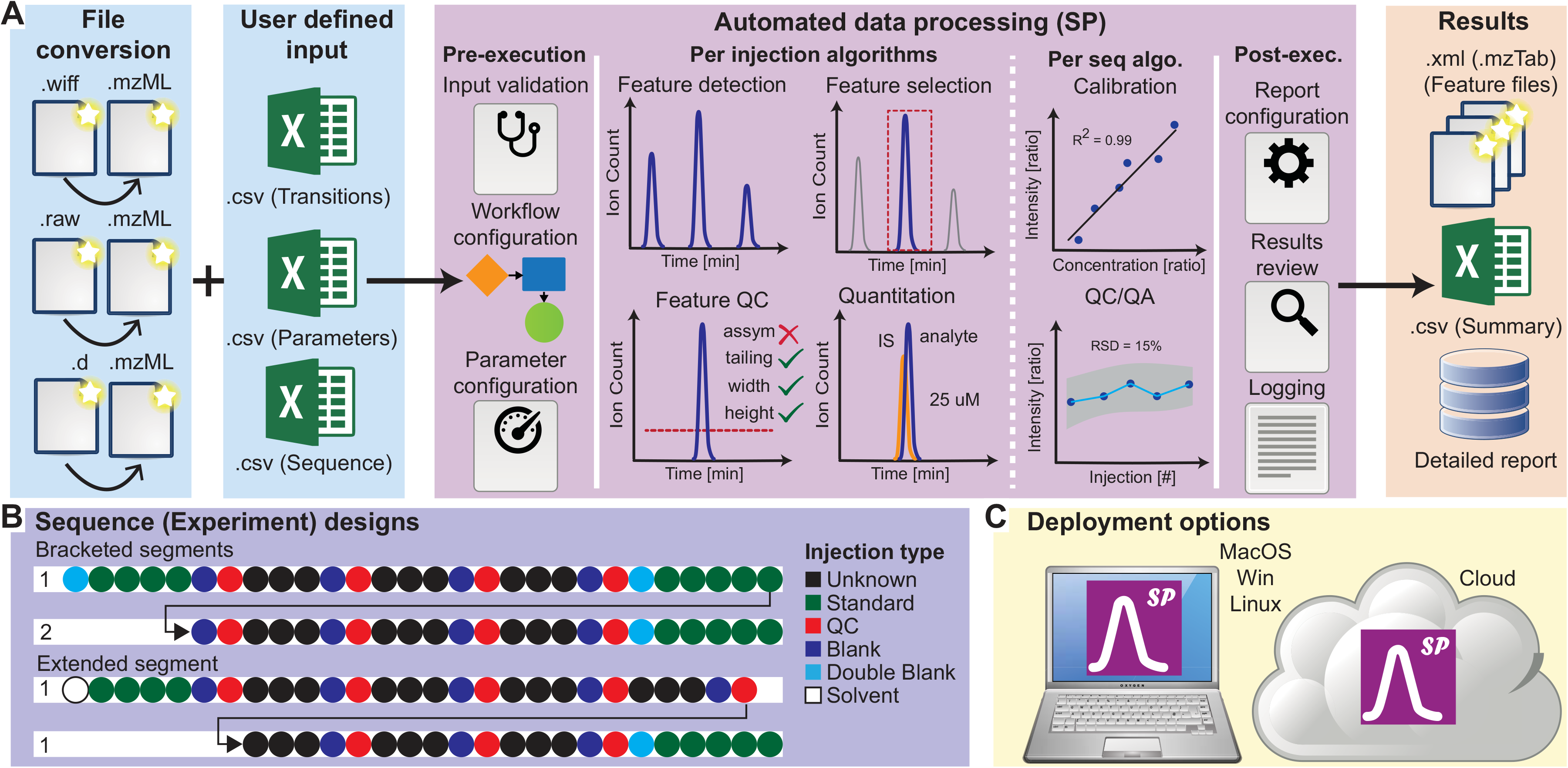
SmartPeak encapsulates advanced algorithms to enable fast, accurate, and automated processing of CE-, GC- and LC-MS(/MS) data, and HPLC data for targeted, semi-targeted, and untargeted metabolomics, lipidomics, fluxomics, and proteomics experiments. A) Custom data mover scripts are used for moving data from an instrument computer to a file store server (not shown). MSConvert from ProteoWizard is used for converting from proprietary file formats to open-source formats. SmartPeak then takes the raw open-source data along with user supplied input and processes the data to generate individual processed data files for each injection in the sequence that can be consumed by other open-source tools along with a formatted summary report and detailed report that is suitable for consumption by LIMS or other database systems. SmartPeak wraps underlying algorithms from OpenMS into a user-centric application by providing features such as input checking, workflow configuration, and parameter configuration prior to executing a data processing workflow on a user-defined sequence along with features such as report configuration, results review, and logging after executing a data processing workflow but prior to reporting the results. The application provides parallelism at both the injection processing level and at the sequence segment processing level that scales to the number of nodes made available prior to launching the data processing workflow. B) Supported sequences or experimental designs include bracketed and extended segments to allow for the application of calculations derived from Standard, Blank, and/or QC samples to either be applied within each segment (bracketed) or calculated once and applied to the whole sequence (extended). Supported injection types include Unknowns (normal samples of unknown concentration), Standards (samples of known concentration that are used for the creation of the calibration curve), QCs (samples of known concentration or of unknown concentration but known composition e.g., pooled Unknowns that are used to check the accuracy of detection and quantification; see ^22^ for an extended discussion of possible QCs), Blanks (samples containing the internal standard compounds, if used, but no analytes, and which have been through the normal sample preparation procedure), Double Blanks (samples with neither internal standards nor analytes), and Solvents (double blanks that have not been through the normal sample work-up procedure). C) SmartPeak can be deployed on a user’s computer for e.g., assay development where greater levels of visualization and debugging are required or in the cloud for e.g., full automation of validated and data intensive workflows.

## Baseline detection and peak integration

Accurate and reproducible quantitation by SIM, SRM, PRM, or HPLC relies heavily upon precise baseline detection and integration of acquired chromatographic peaks (Fig S1). SmartPeak provides users with several options for baseline detection and peak integration. While these methods result in accurate calculations of peak heights and peak areas when peaks are well acquired (e.g., high signal to noise ratio with limited peak tailing), it may be the case that peaks are either saturated (i.e., above the limits of detection) (Fig 2D) or cutoff (Fig 2E). This results in an underestimation of the actual peak height and area that leads to an underestimation of the amount of metabolite or peptide in the sample. A machine learning algorithm was employed that models the peak shape to an exponentially modified gaussian (EMG) curve (see methods for algorithm details). The EMG distribution has been used previously as a model of chromatographic peaks due to its ability to depict chromatographic abnormalities such as shouldering and tailing that are often seen ^15,16^. By fitting each peak to an EMG distribution, the peak height and area of saturated and cutoff peaks can be interpolated closer to their true values (Fig 2G-H, Table S2). Accurate recapitulation of poorly acquired peaks prevents analysts from having to re-prepare, re-inject, and re-analyze the sample, which leads to more accurate results and time saved. Accurate recapitulation of poorly acquired peaks also allows for more flexibility in method development by providing robustness to retention time shifts and variations in on column sample amount.

**Fig. 2.**
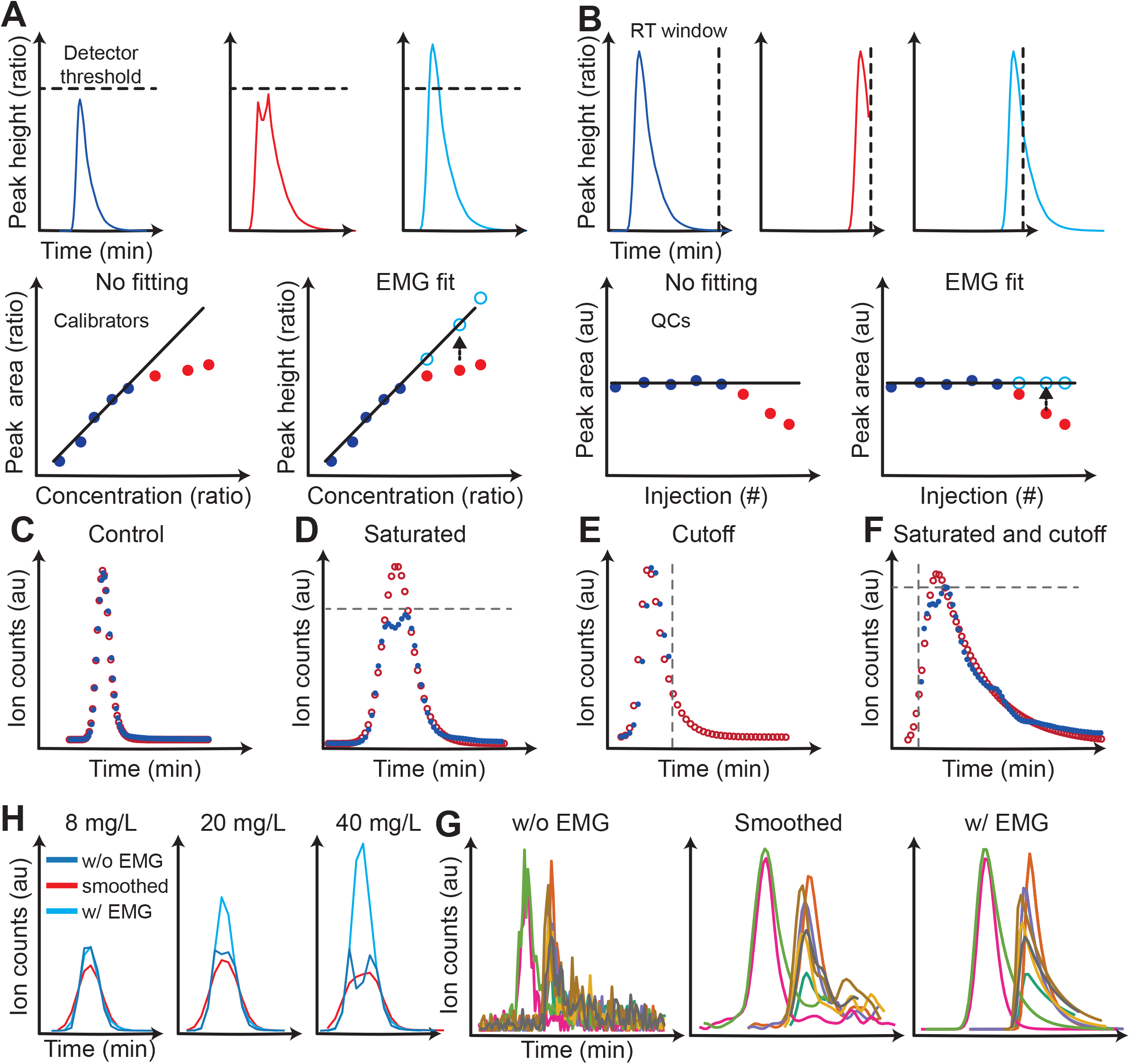
EMG peak interpolation. Applications of EMG peak interpolation are shown for A) extending the upper limit of quantitation (ULOQ) for calibrators by correcting for detector saturation and B) improving the accuracy and precision of quality control (QCs) samples by correcting for cut-off peaks. The red points and lines represent uncorrected data and the light blue points and lines represent EMG interpolated data. Examples of using the EMG peak interpolation method on real chromatographic data for C) an ideal peak, D) a saturated peak, E) a cutoff peak, and F) a saturated and cutoff peak are shown. The blue points represent uncorrected data and the red points represent EMG interpolated data. G) Example application of the EMG algorithm on Standard samples from the BloodProject (Table S3). The EMG algorithm increased the ULOQ from 8 mg/L to 40 mg/L for riboflavin. Dark blue is the original chromatogram. Red is the gaussian smoothed chromatogram. Light blue is the EMG fitted chromatograms. H) Example application of the EMG algorithm to correct for noise in chromatograms for the metabolite UTP in QC samples taken from BloodProject (Table S3). The peak integration of EMG fitted peaks led to a decrease in RSD for UTP from 0.89 to 0.78. The original, smoothed, and EMG fitted chromatograms are shown.

The EMG fitting algorithm is demonstrated on the saturated peaks for calibration curves (Fig 2G) and noisy chromatographic shapes in QC biological replicas of human plasma (Fig 2H). The overall distribution of the relative standard deviation values for the QC compounds was not significantly affected (Fig S7). Thus, using EMG fit for the peak shape correction improves the quantitation accuracy of compounds that were poorly acquired (i.e., saturated, cutoff, or at the limits of detection) while not harming the overall quantitative performance. It should be noted that the EMG curve modeling increases the runtime of the standard SmartPeak workflow by several times.

## Absolute quantitation and automated calibration fitting

The relationship between the amount of metabolite or peptide in a sample and the instrument detector output is established by a calibration curve. For MS detectors, isotope dilution mass spectrometry (IDMS) is routinely used owing to its ability to account for variability in sample preparation, metabolite or peptide degradation, and ion suppression ^17^. IDMS involves spiking in a known amount of isotopically labelled identical or similar components (e.g., 13C labeled) into an unknown sample prior to sample workup (Fig. S2). Standards are added at concentrations that span the lower and upper limits of quantitation (L/UOQ) (Fig. S2). A mathematical model (e.g., linear, y=mx+b) is used to fit the known concentrations in each standard to measured peak areas or heights. Linear, Spline, and Piecewise models with various x and y-axis weighting schemes are currently supported. The model is then inverted when calculating the concentration in an unknown sample.

The calibration of hundreds of metabolites or peptides is a time-consuming task. SmartPeak employs an iterative brute force method that automatically optimizes the calibration curve for each metabolite or peptide given a set of criteria, such as maximal model bias for any calibrator level, minimal R-squared, and minimal acceptable number of points used to fit a calibration model (Fig. S2, methods). The algorithm not only provides an unbiased removal of outliers and results in consistently generated calibration models from batch to batch, but greatly reduces the time spent calculating and reviewing calibration curves (see EMG model fitting and re-analysis of published data sets sections). The automated fitting was applied to 6 calibration curves for 116 compounds (Table S2). The absolute change in R-squared after applying the automated fitting was under 0.01 for 90% of the compounds tested, LLOQ got lower for 51% of the compounds and higher for 24%, ULOQ got higher for 75% of the compounds and lower for 16%.

As it is not possible to use IDMS with HPLC detectors, the algorithm also allows the user to either not use an internal standard or designate another internal or spike-in metabolite or peptide as an internal standard. In addition, more advanced quantitation schemes such as bracketed calibrations are also supported (see examples for HPLC data).

## Adaptive feature alignment and selection

Shifts in metabolite or peptide elution times cause major problems for the reproducible assignment of metabolite or peptide IDs across batches of samples. Most commercial or vendor software assign metabolite or peptide IDs based on a retention time window where the most intense peak or the one with the greatest signal to noise ratio is selected (Fig 3 A and B). The use of absolute retention time windows leads to a misidentification of peaks when two isomers with different abundances elute at similar retention times or when shifts in overall elution times occur, such as when a bubble forms in the column or between batches of mobile phase solvent. The shifting of elution time is particularly problematic in capillary electrophoresis (CE), where shifts of up to 10 mins are not uncommon. When similarly eluting isomers or when chromatographic shifts are present, an operator is forced to either update the retention time window on a sample by sample basis or resort to manually identifying the peak on a peak by peak basis. Either approach is tedious, time-consuming, and error-prone.

**Fig. 3.**
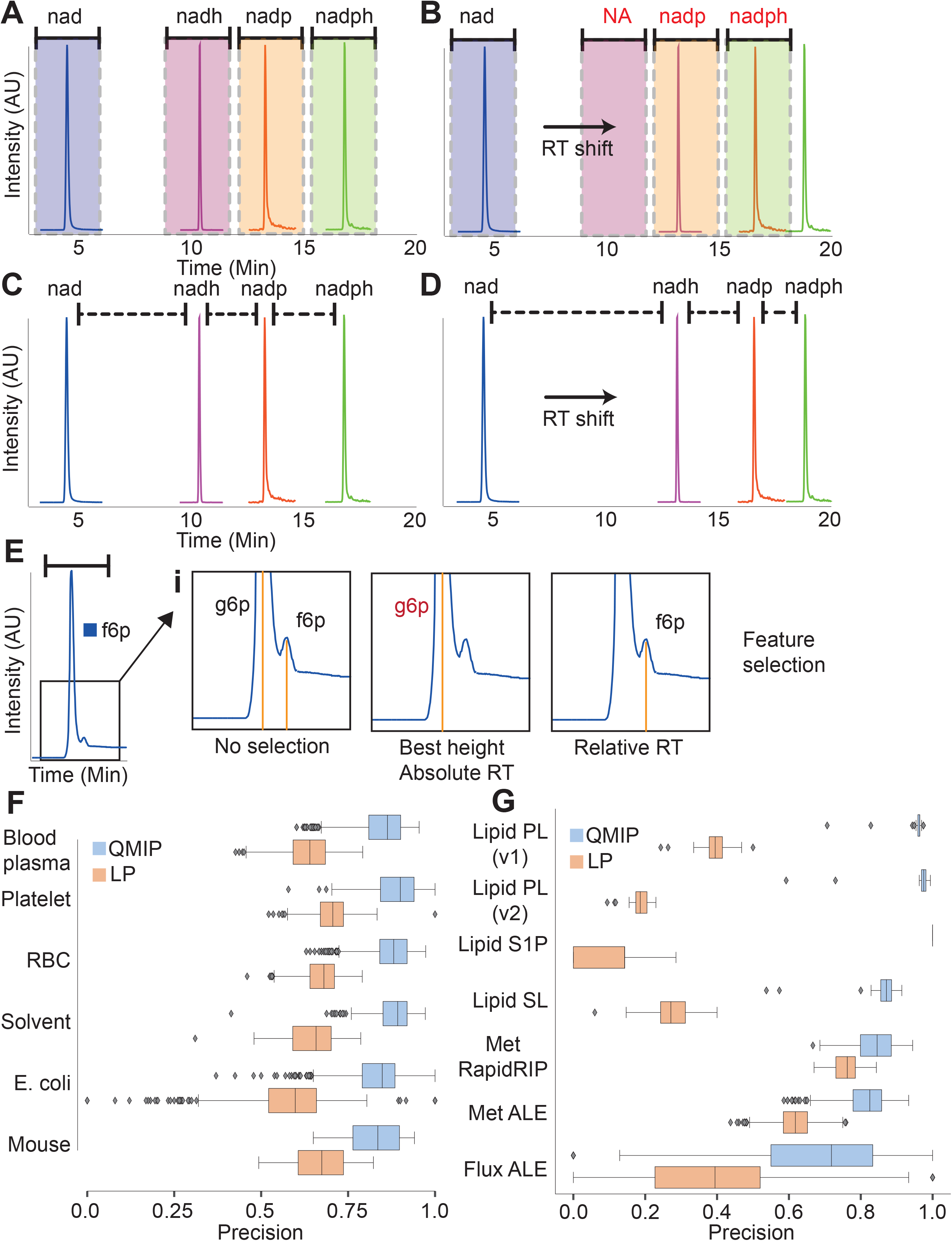
Feature selection algorithms. A) Current feature selection algorithms are based on absolute retention time where a window is set to search for a selected compound. B) Absolute retention time is problematic when retention time shifts occur because the retention time window does not adapt to the retention time shift. This leads to incorrect annotation of compounds that have to be manually corrected by the user. C) Instead, an adaptive feature selection algorithm is implemented that uses quadratic integer programming (QMIP) and relative retention time between components to annotate compounds. D) Relative retention time improves precision of annotating compounds based on retention time by automatically adapting to retention time shifts. E) Relative retention time also improves the annotation of peaks that have large discrepancies in signal intensity. An example chromatogram for the case of glucose-6-phosophate (g6p) and fructose-6-phosphate (f6p) are shown. The larger peak for g6p is often selected over the smaller peak for f6p when using absolute retention time as opposed to relative retention time. See methods further details on the implementation of both absolute and relative retention time feature selection algorithms. F) The precision of automated feature selection tested on metabolomics in different matrices. The boxplot shows the distribution characteristics of a set of samples from the datasets *BloodProject, ALEWt, Fermentors, Mouse* (see Table S3 for datasets descriptions) grouped by the matrix. G) The precision of automated feature selection for reanalysed datasets. The boxplot shows the distribution characteristics of a set of samples grouped by the dataset. The lipidomics samples are additionally grouped by the analytical method: PL (phospholipids, diacylglycerols and cholesteryl esters), S1P (sphingosine 1-phosphate), SL (sphingolipids). The PL method was additionally analysed using another version of the transition list where the split peaks are defined as separate transitions (see Table S1 for the full list of transitions).

As an alternative, retention time calibration components can be spiked into the sample that can be used to automatically adjust the retention time windows. However, this still neglects the case of closely eluting isomers that have overlapping retention time windows (Fig 3E). Instead, we employ an optimization-based approach that assigns metabolite or peptide IDs based on relative retention time (Fig 3 C and D, see methods for algorithm details). The algorithm is formulated as a quadratic mixed integer programming (QMIP) problem that automatically aligns all metabolites or peptides within a sample based on their nearest neighbor distance. Multiple neighbors of each metabolite or peptide can be used, providing robustness and accuracy even when an extensive number of metabolites or peptides are missing in the sample. We find that the algorithm provides superior performance to absolute retention time-based assignment methods (Fig 3 F and H). In particular, isomers with similar retention times but drastically different peak heights are correctly assigned (Fig 3E).

## Flexible and automated feature filtering and quality control

Essential to any automated workflow is the inclusion of quality control and quality assurance (QC/QA) measures that alert users when an algorithm is inaccurate due to an abnormality in e.g. the analytical instrumentation or a sample. Use cases for QC/QA methods span from alerting a technician when a system suitability test (SST) fails due to e.g. fouling of a column, to alerting a bioinformatician when the reproducibility of biological or technical replicates falls below previously defined acceptance criteria. QC/QA methods that are themselves fully automated and systematically applied become absolutely essential when working in Big Data regimes where manual checking of individual samples can only be applied selectively. To this end, we provide a host of customizable filtering, quality control and alerting criteria that can be adapted on a per-method basis (Fig S3). The criteria can be as simple as alerting the user when a metabolite or peptide is outside the range of quantitation or when a peak displays tailing beyond a certain threshold, to more complicated criteria such as when a transition exceeds a predetermined ion ratio (see Methods for an enumeration of all quality control criteria). The information from the QC samples (e.g., pooled biological samples) and Blank samples (i.e., processed samples without analytes) can also be used to flag features that are found to be above a certain level of variability in the QCs or below the level of background interference found in Blank samples. In order to aid the user in developing the initial quality control thresholds, methods are provided to estimate initial values based on Standard, QC and Blank samples. Each of the user-specified criteria is then weighted and combined into a single score that can be automatically filtered on during downstream analysis.

## Performance validation on previously analysed data

To validate the performance of the algorithms described above, reanalysis of several datasets was performed (see Table S3 for the datasets descriptions). We performed two types of analysis: the validation of feature selection algorithm on all the datasets (Fig 3) and automated peak area, peak height and concentration calculation on the following datasets: 1) The metabolomics and fluxomics data for an adaptive laboratory evolution (ALE) experiment with evolved and unevolved *E. coli* strains ^3^; 2) The metabolomics data for a rapid 5 minutes RIP-LC-MS/MS assay for several industrial *E. coli* strains (RapidRIP) ^18^; 3) The lipidomics data for the effect of glucocorticoid exposure in canine plasma ^19^. The last dataset comprised data from different analytical methods, which we processed separately. Pearson correlations between published and calculated peak areas, peak heights, and concentrations are reported in Table n. For most of the experiments the peak area and concentrations correlations were higher than 0.85. The QMIP feature selection algorithm was found to consistently perform better than the linear programming LP algorithm (see Methods for the algorithms description), as determined by the feature selection precision (Fig 2F), peak area and concentration correlations (Table 2). More detailed results are presented in Figures S4 and S5.

**Table. 2.** Correlations of predicted and published values for reanalysed datasets. ‘n’ is the total number of analysed peaks in all the samples.

| Dataset | Peak area, QMIP | Peak area, LP | Concentration, QMIP | Concentration, LP |
| --- | --- | --- | --- | --- |
| Lipidomics, PL (v1) | 0.99<br>n = 8373 | 0.99<br>n = 3354 | 0.97<br>n = 3917 | 0.96<br>n = 1594 |
| Lipidomics, PL (v2) | 0.99<br>n = 6973 | 0.99<br>n = 1348 | 0.99<br>n = 3673 | 0.98<br>n = 693 |
| Lipidomics, S1P | 0.87<br>n = 216 | 0.77<br>n = 31 | 0.99<br>n = 140 | 0.99<br>n = 4 |
| Lipidomics, SL | 0.91<br>n = 2556 | 0.94<br>n = 809 | 0.88<br>n = 614 | 0.70<br>n = 218 |
| Metabolomics, RapidRIP | 0.90<br>n = 25178 | 0.98<br>n = 6396 | 0.85<br>n = 2931 | 0.95<br>n = 435 |
| Metabolomics, ALE | 0.95<br>n = 147888 | 0.96<br>n = 80544 | 0.39 / 0.87*<br>n = 4249 | 0.09 / 0.92*<br>n = 1169 |
| Dataset | Peak area, QMIP | Peak area, LP | Peak height, QMIP | Peak height, LP |
| Fluxomics, ALE | 0.92<br>n = 13010 | 0.93<br>n = 6818 | 0.40 / 0.98**<br>n = 13010 | 0.43 / 0.98**<br>n = 6818 |

\*a correlation calculated without the outlier compound (orn.orn\_1.Light)
\*\*a correlation calculated without the outlier compound group (fdp)

The transition list for phospholipids, diacylglycerols and cholesteryl esters was modified to include additional transitions (Table S1) that were previously included into the dataset as retention time qualifiers only. The second transition list specified each of the split peaks as a separate transition and allowed the QMIP algorithm to take advantage of the relative retention time differences to identify and integrate each of the different split peaks more accurately, which leads to increased precision and recall (Lipid, PL (v1) vs Lipid, PL (v2) on Fig 3G, Methods, Fig S5). This demonstrates that the performance of peak detection and integration using the QMIP algorithm increases as more transitions are added to the method.

The Pearson correlation coefficients for the peak areas and concentrations in Table 2 do not always match (for Lipidomics, S1P the concentration correlation is close to 1 but the peak area correlation is <0.9; for Metabolomics, ALE the peak area correlation is 0.95 but the concentration correlation is <0.9). There are several reasons for this. First, if noisy chromatograms are present in the dataset, the peak area correlation can decrease because of the systematic difference in how the peak baseline and peak start/stop positions are defined for this subset of chromatograms. Some examples of chromatograms that can cause such discrepancies are shown in Fig S4. This may not affect the peak areas ratios, thus still leading to a high correlation in concentrations. Second, differences in the calibration curve optimisation process can lead to decreased concentration correlations despite the well correlated peak area values. And third, the Pearson correlation coefficient itself can be affected by a few outliers. However, we find that the magnitude of these discrepancies between the automated and manual data processing workflows does not affect the biological insights derived from the reanalyzed data (Fig S6) using the example of the canine lipidomics study.

## Scalability to Big Data and cloud infrastructures

Users can access the algorithms described above directly from C++ or Python via the PyOpenMS ^20^ wrapper, where they can be combined into custom scripts that can be run on any computational infrastructure. To enable increased reproducibility and ease of access in regulated environments, a GUI was made that allows the user to easily load all raw data files and input data files, run a consistency check on all provided input prior to analysis, configure, monitor, and run workflows, visually inspect the results, debug problematic samples or features through visual and algorithmic procedures, and export the results to .csv format. The application automatically logs all user commands and application messages for error checking and audit logging. Importantly, the workflows can be run utilizing multiple threads where each thread applies a workflow to a different sample so that large datasets can be processed in parallel. The GUI is written in C++ using the ImGui framework (https://github.com/ocornut/imgui/) along with various thread and data caching strategies to maximize performance. Example workflows for metabolomics and fluxomics applications using LC-MS/MS, HPLC, and GC-MS are provided with the source code. The source code for SmartPeak can be found at https://github.com/dmccloskey/SmartPeak2, and the latest stable builds of the GUI can be found at https://github.com/dmccloskey/SmartPeak2.

## Conclusion

We have found that SmartPeak can reduce what was once considered a multi-week data process workload using commercial and vendor-specific software into a day with a trained user. For example, the 5332 samples included here required approximately 5332 hours (~ 60 minutes per sample) to manually process; with SmartPeak, the same 5332 samples required 11.0 hours (~1 min per sample per thread using 8 threads of an i7-6820HQ 2.7 GHz processor with 16 GB of RAM) to automatically process. This equates to an over 100 fold reduction in processing time using a circa 2018 laptop with the added benefit of improving quality and reproducibility. The algorithms described above raise the bar for accurate, fast, and automated data processing in targeted and/or quantitative metabolomics, lipidomics, and fluxomics applications across a multitude of chromatography and mass spectrometry instrumentation found in the analytical chemistry field. As a part of the OpenMS toolkit, the algorithms are easily extensible and modifiable for future workflows.

## Methods

### Peak integration methods

Several peak integration methods and modes for specifying the baseline of the peak are available to the user. The different peak integration methods and modes for specifying the baseline of the peak are graphically depicted in Fig S1. Peak integration methods include Intensity Sum where the sum of all peak intensities are used to define the peak area; Trapezoid where the trapezoid rule is used to calculate the peak area; and Simpson where the simpson’s formula is used to calculate the peak area. Modes of specifying the baseline of the peak include “Base To Base” which extends a straight line from the peak start to the peak end; “Vertical division min” where a horizontal line is drawn from the lowest peak start or end point to a vertical line that is drawn from the highest peak start or end point; and “Vertical division max” where a horizontal line is drawn from the highest peak start or end point to the edge of the peak.

### EMG peak interpolation

Peaks were modeled as exponentially modified gaussians (EMG) following the equations derived in ref ^16^. In brief, for a given z value, a point on the EMG curve was calculated as follows::

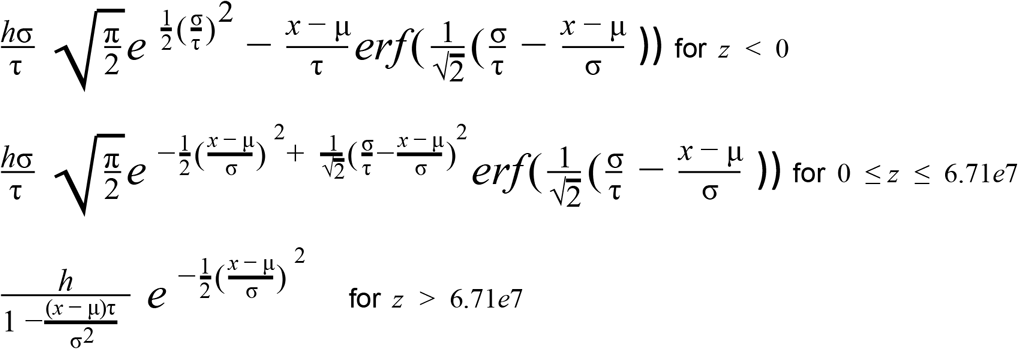

where *x* is the position (either time or m/z), *σ* is the Gaussian height or amplitude, a is the Gaussian standard deviation, μ is the position of the Gaussian mean, and τ is the exponent relaxation time parameter used to modify the Gaussian. *z* is a constant calculated using the following formula:

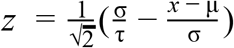

Stochastic gradient descent (SGD) using the iProp+ algorithm ^21^ was used to fit the h, sigma, mu and tau parameters using a mean squared error (MSE) loss function. The MSE loss was computed from the known intensity of the training data point and the estimated intensity given the data point position and current EMG parameters. The initial value for h was set as the most intense peak value. The initial value for mu was determined by estimating the most probable position for the peak apex using a heuristic that provides robustness to spurious points or random fluctuations in the detector sampling that would result in an estimation of mu that is far from the actual peak apex. Candidate points for the initial value of mu were selected by taking the average of the position of all points less than or equal to 60, 65, 70, 75, 80, and 85% of the most intense data point. Initial values for sigma and tau were then determined by multiplying mu by empirically determined constants that gave optimal results from several example data sets. iProp+ solver parameters were determined empirically based on optimal settings from several example data sets.

Training data for modeling individual chromatographic peaks were extracted from regions of the chromatogram that were not saturated (i.e., where the intensity was above the instrument detector limits). A heuristic for determining non-saturated data points was employed that checked the derivative of all points with an intensity greater than 80% of the most intense peak. Given a continuous region of points that are greater than 80% of the most intense peak, starting from the left, points were added to the training data until the derivative reached a value greater than 0.3 or the ratio of the current derivative and the previous derivative exceeded 0.6. Starting from the right, points were added using the same derivative thresholds. Derivatives were estimated using the finite difference method between the current point and the previous point. Thresholds used in the described heuristic above were determined empirically based on optimal settings from several example data sets.

Once the optimal EMG parameters were found, they were then used to recompute the intensity of each peak based on the original positions. For the case of cut-off peaks, the original chromatographic positions were extended either left or right until both the starting left and ending right positions reached the same intensity values. Several examples on test data are given in Fig 2

### Automated calibration curve fitting

Given a mathematical model (e.g., linear model y = mx + b) that relates the concentration ratio, x, to peak area or height ratio, y, with optimal weighting (e.g., inverse 1/x), a small number (i.e., less than 100) of samples with known x and y values can be iteratively searched efficiently to generate the best model parameters (i.e., m and b in the case of a linear model) that meet a given set of criteria (Fig S2). For the purposes of calibration curve fitting, the criteria are model bias for any calibrator level, the minimal R-squared value of the model, and the number of points used to fit the model.

Model bias, b, was calculated using the following formula: b = fabs(actual_concentration – calculated_concentration)/actual_concentration*100;

Model R-squared, r2, was calculated as the square of the Pearson’s correlation coefficient.

The algorithm starts by calculating the model bias and model R-squared for all points. If the criteria for minimal model bias and minimal model R-squared are met, the algorithm returns the fitted model parameters. If not, the algorithm removes a single point that is determined to be the largest outlier using either the jackknife method or residual method.

The jackknife outlier detection method selects candidate outliers by iteratively removing a single data point and calculating the R-squared value. The removed data point that leads to the highest R-squared value is selected as the most probable outlier.

The residual outlier detection method selects candidate outliers according to the data point with the largest residual error (i.e., bias).

The algorithm continues to iteratively refit the model and remove candidate outliers until the minimal model bias and minimal model R-squared criteria are satisfied or there are fewer remaining points to fit the model to than what was specified as the minimum.

### Feature selection

#### The peak selection optimization algorithm

Peaks were detected using the MRM feature finder algorithm provided with OpenMS ^14^ library. Peaks were then selected using an optimization algorithm that accounted for peak quality and nearest neighbor retention time difference (Fig 2). The quadratic mixed-integer problem formulation was the following:

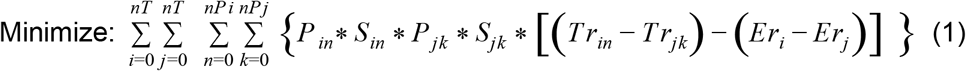

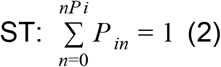

and

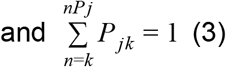

Where:

nT is the number of transition

nP is the number of peaks

i and j are indices over all transitions

n and k are indices over all peaks per transition

*P* ∈*Z*, ∀ ∈*I* P is a binary variable indicating Peak n or k for Transition i or j is included or excluded, respectively

Tr is the retention time for Peak n or k for Transition i or j.

Er is the expected retention time for Transition i or j.

W is an optional weighting factor that scales inversely according to the distance of *Tr*_*in*_ from *Tr*_*jk*_

S is an optional and user defined score for Peak n or k for Transition i or j.

S was typically calculated as follows:

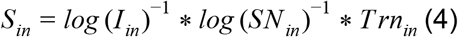

Where I is the intensity, SN is the signal to noise ratio, and Trn is the absolute difference for Peak n or K for Transition i or j and the expected retention time.

The quadratic term can be linearized as follows:

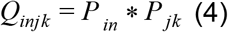

Where

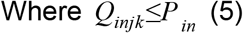

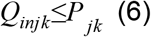

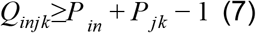

And *Q*∈*Z*, 0≤*Q*≤l Q is a continuous variable

The absolute value term can be linearized as follows:

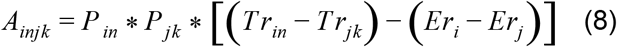

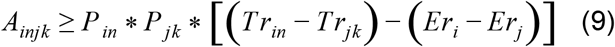

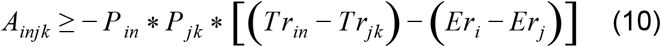

And *A*∈*Z*, − ∞≤*A*≤∞ A is a continuous variable

The linearized problem is thus:

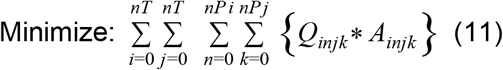

ST: equations (2), (3), (5), (6), (7), (9), (10)

An LC-MS/MS run with over 300 transitions (over 100 compounds) can be loaded, picked, scored, filtered, selected, and quality controlled in less than 90 seconds on a typical laptop (Intel Core i7 6820HQ CPU @2.70 GHz with 16.0 GB of RAM).

#### The control peak selection algorithm

Peaks were first filtered based on a minimum peak height threshold (100 ion counts), log signal to noise threshold (0.5), retention time difference between the peak and expected retention time (0.5 min). The largest peak for each transition was then selected. The MILP formulation for peak selection was the following:

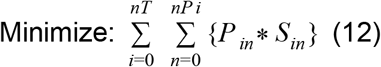

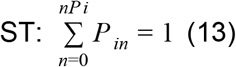

Where S is the score. The score was equivalent to the peak intensity.

#### The performance estimation

The metrics for comparing the feature selection algorithms performance are precision and recall, calculated as:

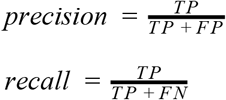

where TP is a number of true positives, FP is a number of false positives and FN is a number of false negatives compared to a reference feature set. A feature is considered a true positive if the retention time (RT) of an automatically selected feature and the RT of a reference feature (i.e. a manually selected one) differ less than the assigned parameter value (3 seconds for the datasets listed in Fig 2). A feature is considered a false positive if the retention time differs more. A feature is considered a false negative if it is a part of a reference set but was not selected with the algortihm.

### Feature quality control metrics

Feature quality control metrics for each injection at the level of the transition/component group and transition/component are made available to the user for building customized filtering or checking criteria on. At the transition group level, the user is able to specify thresholds for retention time, total ion intensity, overall transition quality (i.e., sum of individual transition qualities), the number of heavy (i.e., isotopically labeled with e.g., 13C) or light components, the number of detecting, quantifying, and identifying transitions, and ion ratio (i.e., the intensity ratio between a quantifying transition and a qualifying transition) as well as custom metrics that the user can define. At the transition level, the user is able to specify thresholds for retention time, ion intensity, transition quality, and custom metrics such as the limits of quantitation and various statistical metrics that can be used to describe the shape and integration quality of a chromatographic peak (Fig S3). Specifically, the following metrics are made available to the user for statistically describing the peak shape and integration quality:

‘width_at_5’: The width of the peak at 5% the peak’s height

‘width_at_10’: The width of the peak at 10% the peak’s height.

‘width_at_50’: The width of the peak at 50% the peak’s height.

‘start_position_at_5’ and ‘end_position_at_5’: The start and end positions at which the intensity is 5% the peak’s height.

‘start_position_at_10’ and ‘end_position_at_10’: The start and end positions at which the intensity is 10% the peak’s height.

‘start_position_at_50’ and ‘end_position_at_50’: The start and end positions at which the intensity is 50% the peak’s height.

‘total_width’: The peak’s total width as defined by the start and end positions.

‘tailing_factor’: The tailing factor is a measure of peak tailing. It is defined as the distance from the front slope of the peak to the back slope divided by twice the distance from the centerline of the peak to the front slope, with all measurements made at 5% of the maximum peak height. The formula for calculating the tailing_factor, Tf, is Tf = W0.05/2a where W0.05 is peak width at 5% max peak height, a = min width to peak maximum at 5% max peak height, b = max width to peak maximum at 5% max peak height. A Tf value between 0.9 and 1.2 indicates no peak tailing; a Tf value less than 0.9 indicates peak fronting; and a Tf value greater than 1.2 indicates peak tailing.

‘asymmetry_factor’: The asymmetry factor is a measure of peak tailing. It is defined as the distance from the centerline of the peak to the back slope divided by the distance from the centerline of the peak to the front slope, with all measurements made at 10% of the maximum peak height. The formula for calculating the asymmetry_factor, As, is As = b/a where a is min width to peak maximum at 10% max peak height and b is max width to peak maximum at 10% max peak height.

‘slope_of_baseline’: The slope of the baseline is a measure of slope change and is approximated as the difference in baselines between the peak start and peak end.

‘baseline_delta_2_height’: The change in baseline divided by the height is a way of comparing the influence of the change of baseline on the peak height.

‘points_across_baseline’: The number of points across the baseline

‘points_across_half_height’: The number of points across half the peak’s height.

In addition to the above feature quality control metrics that operate on the metric values of the features themselves, the user is also able to specify thresholds for any of the metrics above for the amount of variability as measured by percent relative standard deviation (%RSD) as measured in QC samples (i.e., pooled biological replicates), and thresholds for minimum acceptable percent background interference as measured by ion intensity for the transition group or transition compared to Blank samples (i.e., matrix matched and processed samples without analytes but with internal standards).

The quality control score at the feature or feature group level is a sum of all passed metrics (i.e., 1 for pass and 0 for fail) that are normalized to the total number of metrics and expressed as a percent.

### Automated feature filtering, flagging, and initial threshold limit estimations

Methods are provided to allow for filtering and flagging based on any of the feature quality control metrics discussed previously. The filtering methods remove transition groups or transitions that fail to pass one or more user defined thresholds, while the flagging methods report failed thresholds without removing the transition or transition group. In addition, the flagging methods build an overall quality score based on the number of passed or failed thresholds that the user can then filter on when performing downstream data analysis tasks. The filtering methods provide means for the user to remove poor quality data systematically, while the flagging methods provide means for the user to score and report the quality of the data.

In order to automate and systematize the process of developing the feature quality control metrics, methods were implemented to estimate user specified quality control metrics based on generated data. The methods include an estimate of the quality control threshold metrics based on the lower and upper ranges of values found in Standard, QC, and/or Blank samples as well as an estimate of the lower and upper limits of quantitation based on the fitted calibration curve. These estimated thresholds can either be used directly by the user to filter or flag features in the same injection sequence or can be further modified (e.g., in order to develop an SOP for the assay) and used in subsequence injection sequences.

## Data availability

All data and parameters used in this study can be found in MetaboLights^23^ repository MTBLS1786.

## Contributions

S.K. and D.M. designed and performed the experiments, wrote the OpenMS algorithms, wrote the SmartPeak application, and wrote the manuscript. P.C. helped write the OpenMS algorithm and helped write the SmartPeak application. T.S., H.R., O.A., O.K. reviewed the OpenMS algorithms and reviewed the manuscript. B.B. and F.T. helped with the lipidomics study data analysis and re-analysis and reviewed the manuscript. L.S. and M.K. helped with the HPLC, GC-MS, and LC-MS/MS examples and reviewed the manuscript. L.N. and M.H. helped design the experiments and reviewed the manuscript.

## Acknowledgements

We would like to thank the entire OpenMS development team for helpful discussions. We would like to thank Rogelio Oseguera from Synthetic Genomics and Robin Palfreyman from the Australian Institute for Biotechnology and Nanotechnology (AIBN) for extensive user testing. This work was funded by the Novo Nordisk Foundation Grant Number NNF10CC1016517. B.B. was funded by the Life Sciences Institute (LSI), National University of Singapore.,

