## Supplemental S01-S07 for "SmartPeak automates targeted and quantitative metabolomics data processing"

### Supplementary material

#### Fig. S1

Peak integration strategies and baseline detection methods.  An example chromatogram from real data with two peaks is shown to illustrate the different peak integration options and baseline detection methods that are available to the user.  i) The peak integration methods available include Intensity Sum, Trapezoid, and Simpson  See methods for a detailed description of each.  ii) The baseline detection methods available include Vertical division (min), Base to base, and Vertical division (max).  See methods for a detailed description of each.

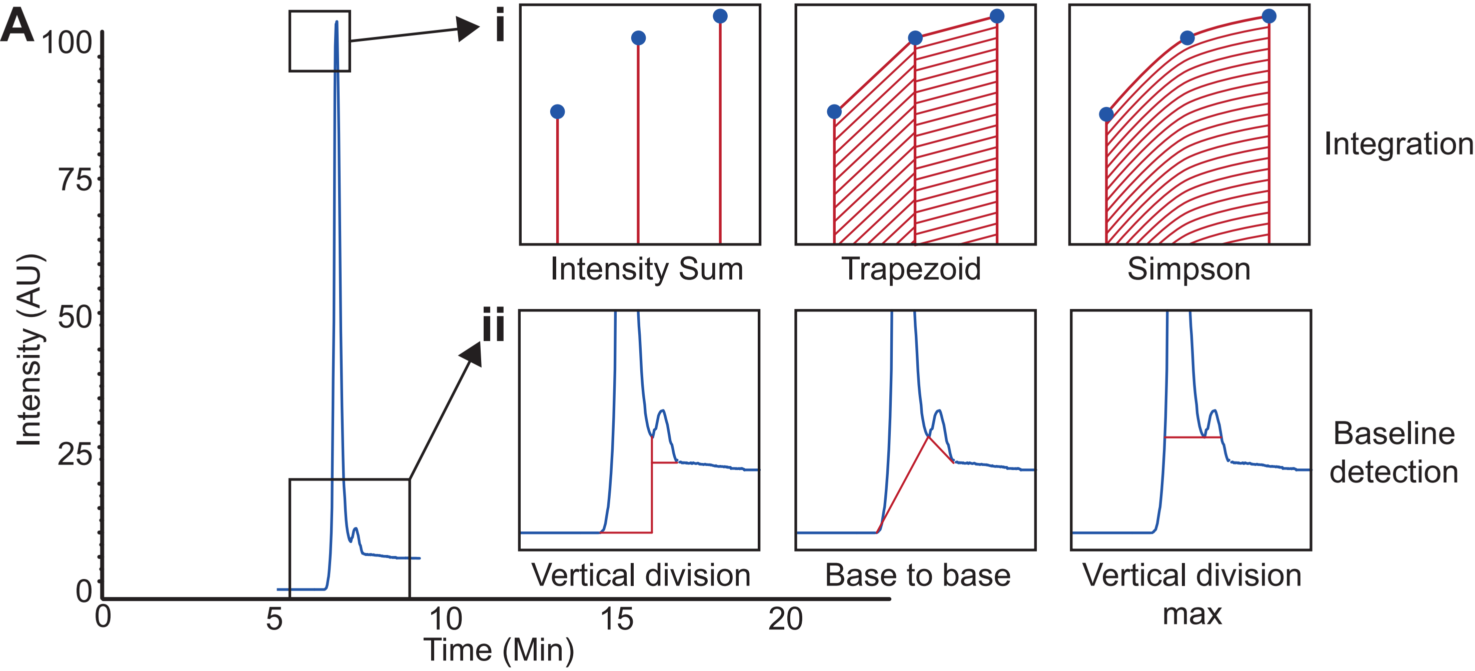

A
i
ii
100
75
50
25
0
Intensity (AU)
0
5
10
15
20
Time (Min)
Intensity Sum
Trapezoid
Simpson
Integration
Vertical division
Base to base
Vertical division max
Baseline detection

### Fig. S2

Automated quantitation and calibration curve fitting. A) A workflow for absolute quantitation using Isotope Dilution Mass Spectrometry (IDMS) is shown. First, Standard samples that consist of a known amount of internal standard (IS) and analyte from stock standards are made. The mass spectrometer can differentiate the internal standard or heavy version (often U13C labeled) of the analyte by the difference in m/z. Second, Standards are created with concentrations that span the linear range of the instrument defined by the upper and low limits of detection (ULOQ and LLOQ). A relationship (e.g., linear as shown) between the concentration ratio and peak area or height of the IS and analyte is then determined. Third, the relationship between the concentration ratio and peak area or height ratio is then used to determine the absolute amount of analyte in Unknown samples that have been spiked with a known amount of IS. B) A brute force calibration curve fitting algorithm is shown that automates the process of determining the relationship between the concentration ratio and peak area or height ratio of Standard samples by iteratively removing outlier points until criteria for the Bias and R2 value are met. See methods for a detailed description of the algorithm.

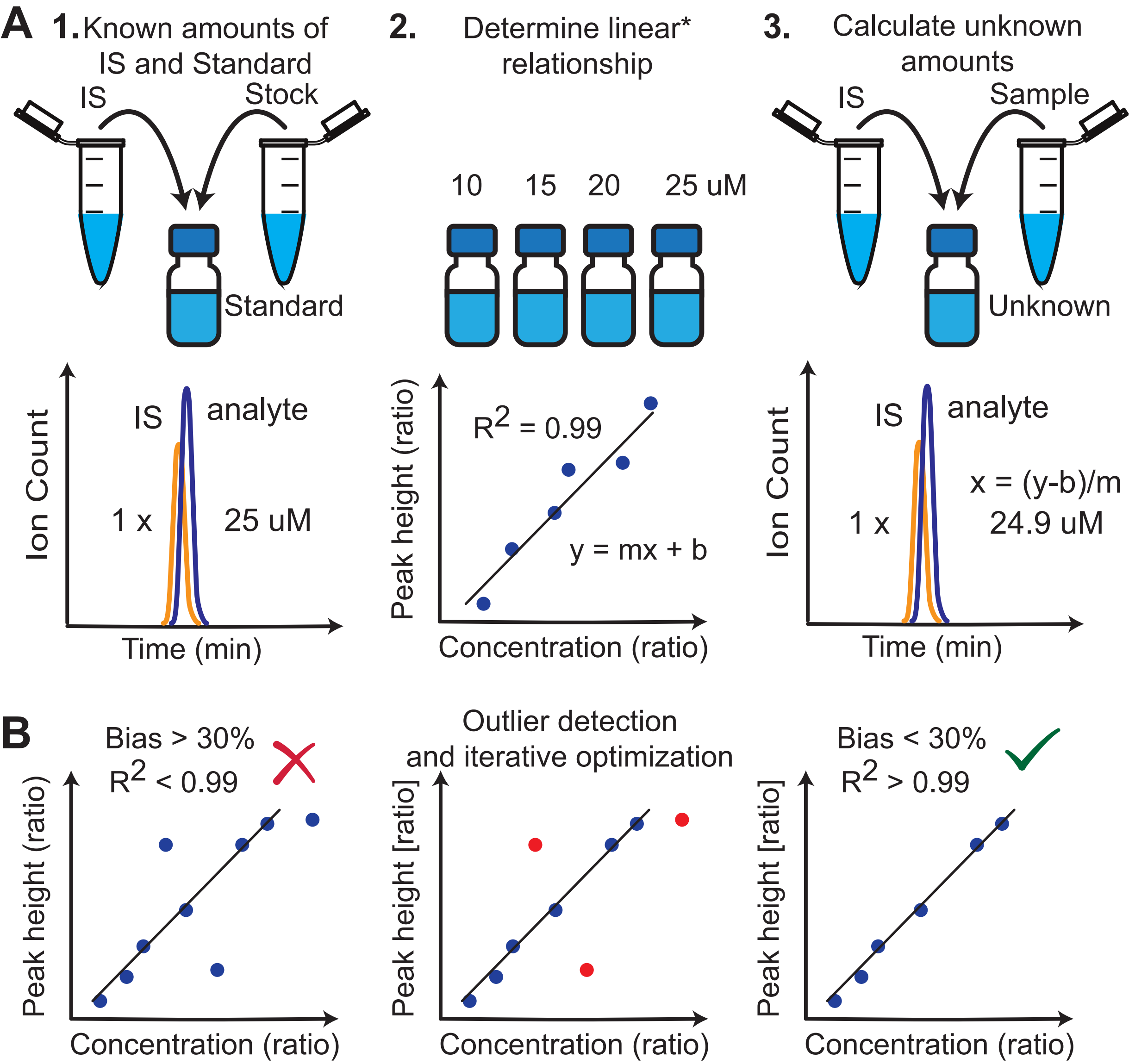
A
1. Known amounts of IS and Standard
IS
Stock
Standard
2. Determine linear* relationship
10
15
20
25 uM
3. Calculate unknown amounts
IS
Sample
Unknown
Ion Count
IS
analyte
1 x
25 uM
Time (min)
Peak height (ratio)
R2 = 0.99
y = mx + b
Concentration (ratio)
Ion Count
IS
analyte
x = (y-b)/m
1 x
24.9 uM
Time (min)
B
Bias > 30%
R2 < 0.99
Peak height (ratio)
Concentration (ratio)
Outlier detection and iterative optimization
Peak height [ratio]
Concentration (ratio)
Bias < 30%
R2 > 0.99
Peak height [ratio]
Concentration (ratio)

### Fig. S3

Quality control. A) Several peak shape metrics (see Methods) are available to the user for flagging and scoring peaks. B) Examples of various peaks from real data with different degrees of peak quality are shown. Common names that would be typically given to classify each peak are given above the chromatograms. Blue points are the raw chromatographic points and red points are the picked peaks. The user is able to set the thresholds for each of the available peak shape metrics in order to flag or filter out low quality peaks in an automated fashion. In the examples shown, quality control thresholds were specified for nine different metrics on a peak by peak basis that included the following: peak height (H), log signal to noise ratio (SN), width at 50% (W50), total width (W), tailing factor (Tf), asymmetry factor (Af), baseline delta to height (dB2H), points across the peak (PH), and points across half height (PH50). The peak quality score based on these thresholds is shown in bold. The failing metrics are shown in italics. C) An example distribution of quality scores (Q) and individual metrics for peak height (H) for Phosphoenolpyruvate (pep) across a batch that includes Standards (green), Blanks (blue), QCs (red), and Unknowns (black). Each circle represents 3 samples. Injection type colors are the same as in Fig 1B.

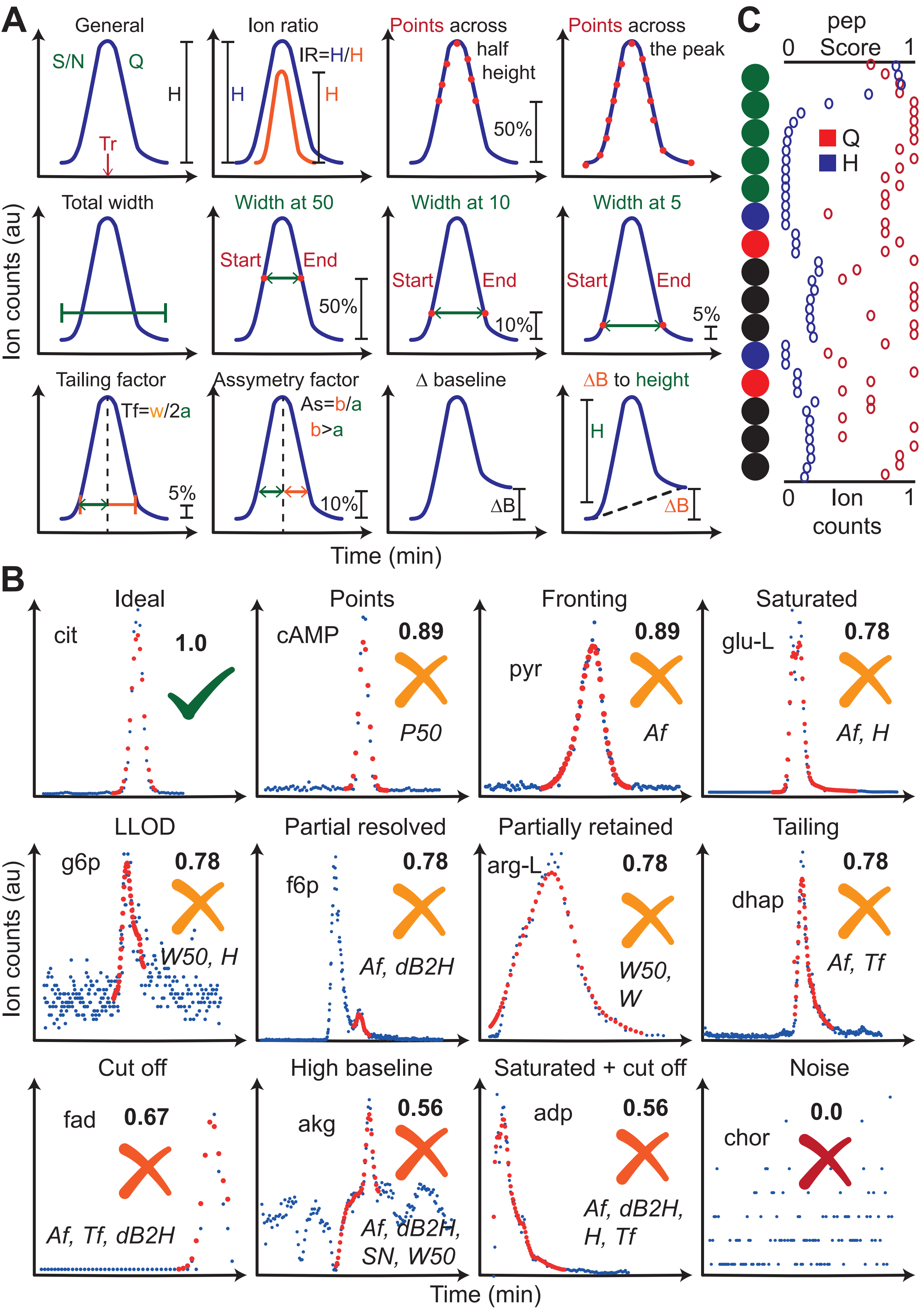
A
General
S/N
Q
H
Tr
Ion ratio
IR=H/H
H
H
Points across half height
50%
Points across the peak
Total width
Width at 50
Start
End
50%
Width at 10
Start
End
10%
Width at 5
Start
End
5%
Tailing factor
Tf=w/2a
5%
Assymetry factor
As=b/a
b>a
10%
Δ baseline
ΔB
ΔB to height
H
ΔB
Ion counts (au)
Time (min)
C
pep Score
0
1
Q
H
0
Ion counts
1
B
Ideal
cit
1.0
Points
cAMP
0.89
P50
Fronting
pyr
0.89
Af
Saturated
glu-L
0.78
Af, H
LLOD
g6p
0.78
W50, H
Partial resolved
f6p
0.78
Af, dB2H
Partially retained
arg-L
0.78
W50, W
Tailing
dhap
0.78
Af, Tf
Cut off
fad
0.67
Af, Tf, dB2H
High baseline
akg
0.56
Af, dB2H, SN, W50
Saturated + cut off
adp
0.56
Af, dB2H, H, Tf
Noise
chor
0.0
Ion counts (au)
Time (min)

### Fig. S4

The scatter plot shows predicted peak areas, height and concentrations against the reference values. The data points from the reanalysed published datasets described in the section “Performance validation on previously analysed data” are grouped by the datasets (Lipidomics, ALE, RapidRIP) and -omics data type (metabolomics, fluxomics, lipidomics). The remaining data points are grouped by the matrix. Some outlier points with large discrepancies between predicted and reference values are additionally annotated with their chromatographic peaks.

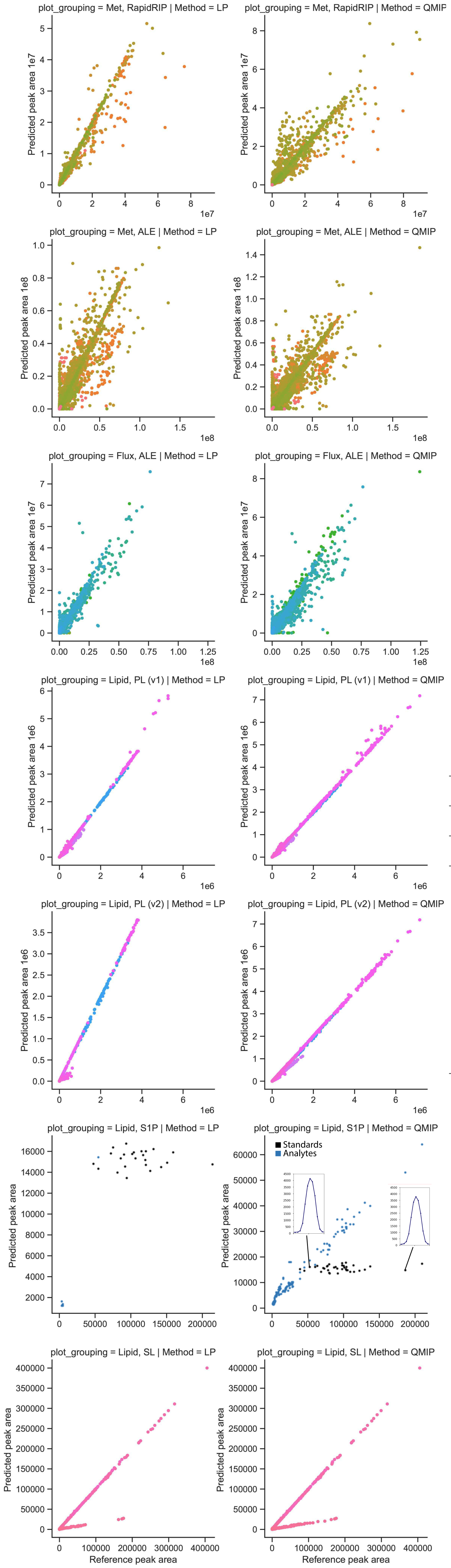
plot_grouping = Met, RapidRIP | Method = LP
plot_grouping = Met, RapidRIP | Method = QMIP
plot_grouping = Met, ALE | Method = LP
plot_grouping = Met, ALE | Method = QMIP
plot_grouping = Flux, ALE | Method = LP
plot_grouping = Flux, ALE | Method = QMIP
plot_grouping = Lipid, PL (v1) | Method = LP
plot_grouping = Lipid, PL (v1) | Method = QMIP
plot_grouping = Lipid, PL (v2) | Method = LP
plot_grouping = Lipid, PL (v2) | Method = QMIP
plot_grouping = Lipid, S1P | Method = LP
plot_grouping = Lipid, S1P | Method = QMIP
plot_grouping = Lipid, SL | Method = LP
plot_grouping = Lipid, SL | Method = QMIP
Predicted peak area 1e7
Predicted peak area 1e8
Predicted peak area 1e6
Predicted peak area
Reference peak area
Standards
Analytes

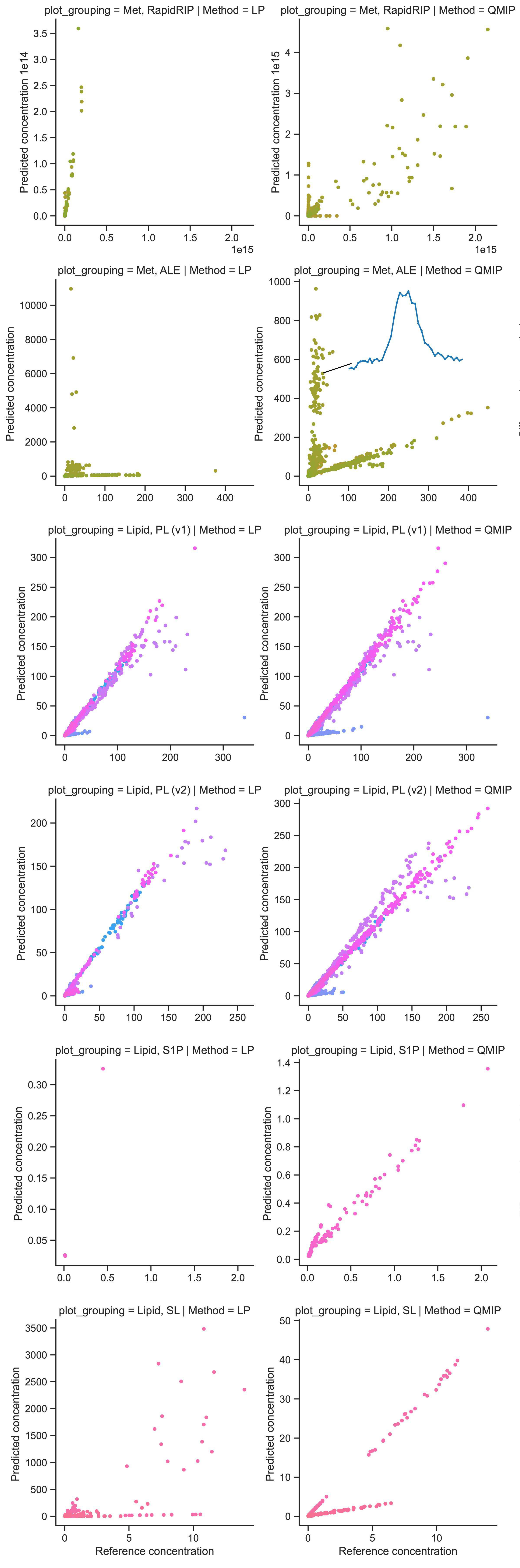
plot_grouping = Met, RapidRIP | Method = LP
plot_grouping = Met, RapidRIP | Method = QMIP
Predicted concentration 1e14
Predicted concentration 1e15
1e15
plot_grouping = Met, ALE | Method = LP
plot_grouping = Met, ALE | Method = QMIP
plot_grouping = Lipid, PL (v1) | Method = LP
plot_grouping = Lipid, PL (v1) | Method = QMIP
plot_grouping = Lipid, PL (v2) | Method = LP
plot_grouping = Lipid, PL (v2) | Method = QMIP
plot_grouping = Lipid, S1P | Method = LP
plot_grouping = Lipid, S1P | Method = QMIP
plot_grouping = Lipid, SL | Method = LP
plot_grouping = Lipid, SL | Method = QMIP
Predicted concentration
Reference concentration

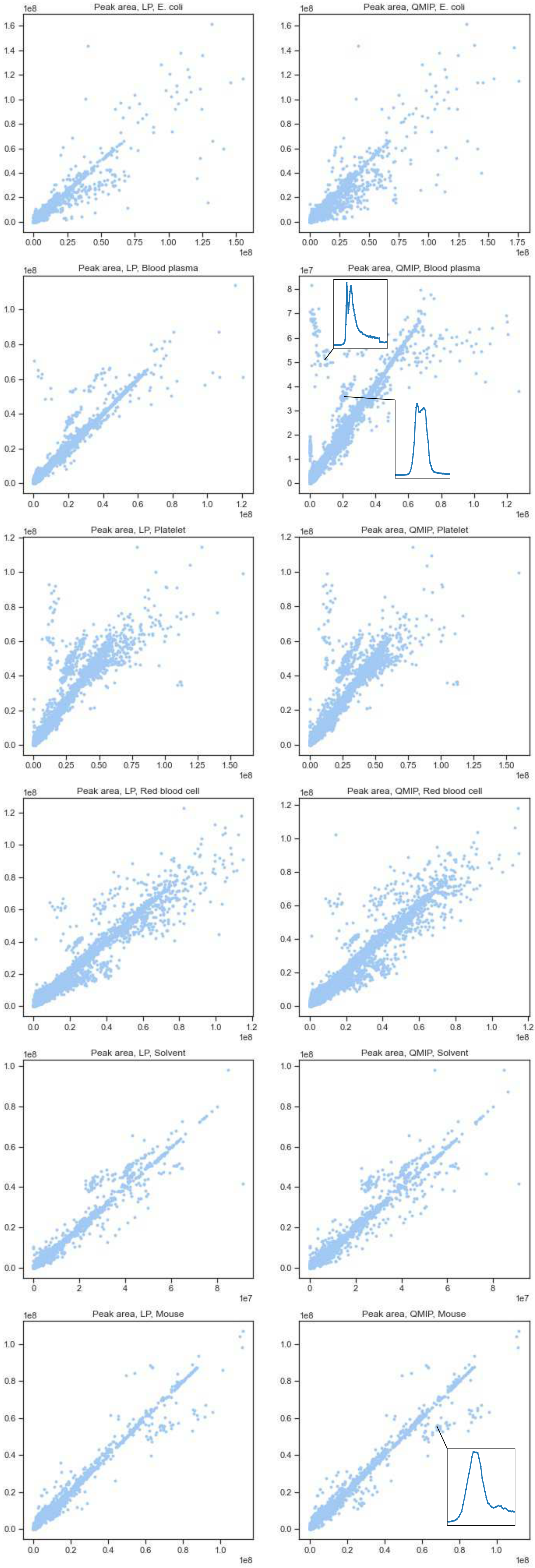
Peak area, LP, E. coli
Peak area, QMIP, E. coli
Peak area, LP, Blood plasma
Peak area, QMIP, Blood plasma
Peak area, LP, Platelet
Peak area, QMIP, Platelet
Peak area, LP, Red blood cell
Peak area, QMIP, Red blood cell
Peak area, LP, Solvent
Peak area, QMIP, Solvent
Peak area, LP, Mouse
Peak area, QMIP, Mouse

### Fig. S5

The recall plots for the analysed datasets (see Methods for the definition). The feature filtering settings have a major effect on the number of features selected.

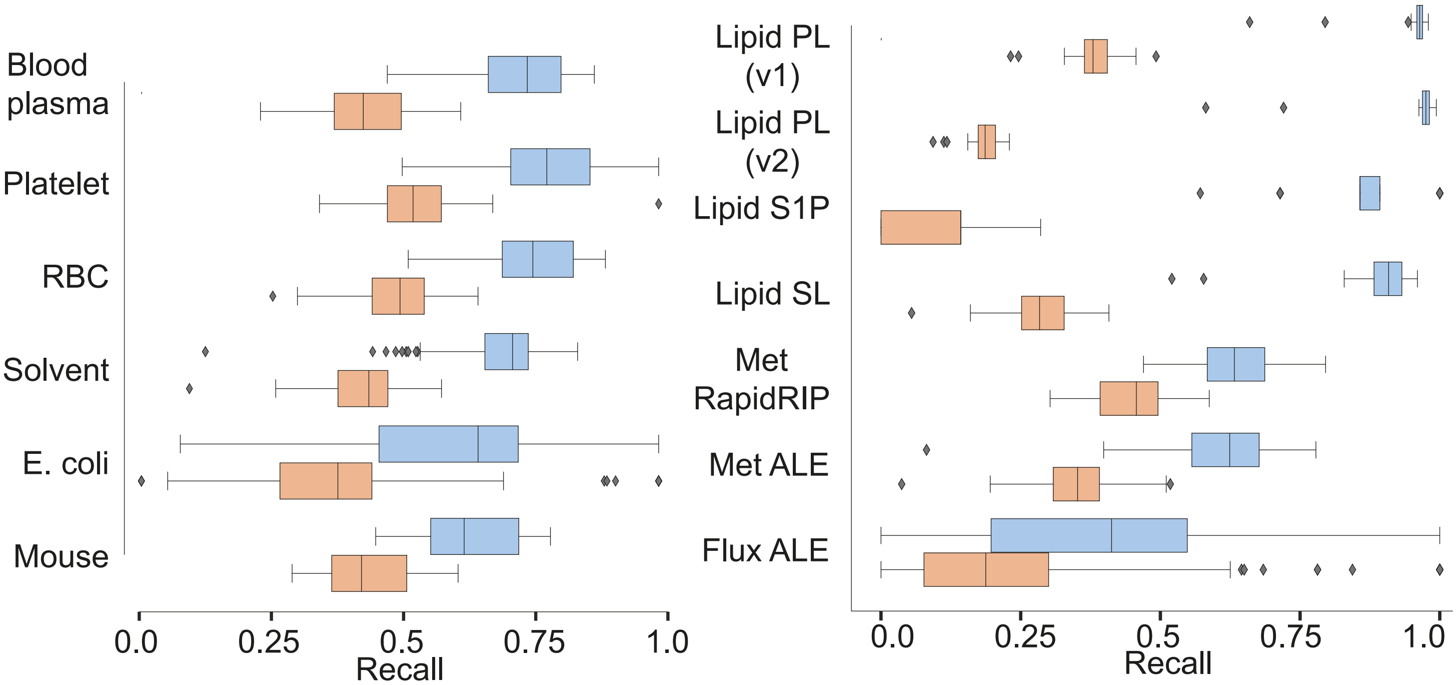
Blood plasma
Platelet
RBC
Solvent
E. coli
Mouse
0.0
0.25
0.5
0.75
1.0
Recall
Lipid PL (v1)
Lipid PL (v2)
Lipid S1P
Lipid SL
Met RapidRIP
Met ALE
Flux ALE
0.0
0.25
0.5
0.75
1.0
Recall

### Fig. S6

The demonstration of the similarity of PCA analysis results for Lipidomics[19] dataset. The first column: original figures from [19]. The second column: the same data analysis performed on the subset of the original data that were reanalysed with SmartPeak. The third column: the SmartPeak reanalysis results. A, B, C and D correspond to c), d), e), f) from Figure 2 from [19].

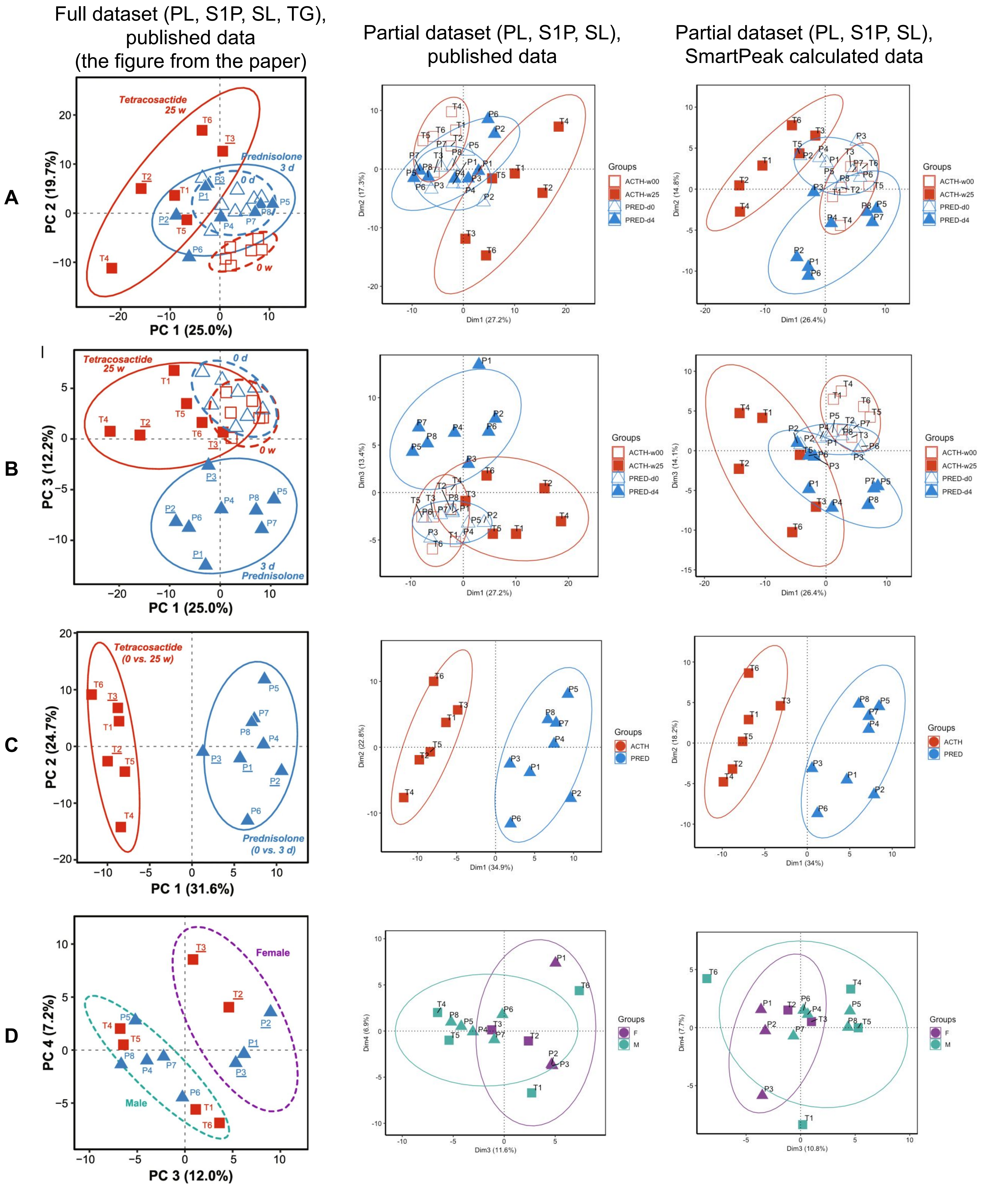
Full dataset (PL, S1P, SL, TG), published data (the figure from the paper)
Partial dataset (PL, S1P, SL), published data
Partial dataset (PL, S1P, SL), SmartPeak calculated data
A
B
C
D
Tetracosactide 25 w
Prednisolone 3 d
0 d
0 w
Tetracosactide (0 vs. 25 w)
Prednisolone (0 vs. 3 d)
Female
Male
PC 1 (25.0%)
PC 2 (19.7%)
PC 3 (12.2%)
PC 1 (31.6%)
PC 2 (24.7%)
PC 3 (12.0%)
PC 4 (7.2%)
Dim1 (27.2%)
Dim2 (17.3%)
Dim3 (13.4%)
Dim1 (34.9%)
Dim2 (22.8%)
Dim3 (11.6%)
Dim4 (6.9%)
Dim1 (26.4%)
Dim2 (14.8%)
Dim3 (14.1%)
Dim1 (34%)
Dim2 (18.2%)
Dim3 (10.8%)
Dim4 (7.7%)
Groups
ACTH-w00
ACTH-w25
PRED-d0
PRED-d4
ACTH
PRED
F
M

### Fig. S7

Distribution of the relative standard deviation values for the peak areas for BloodProject QC samples compounds (reference data, with and without EMG fitting).

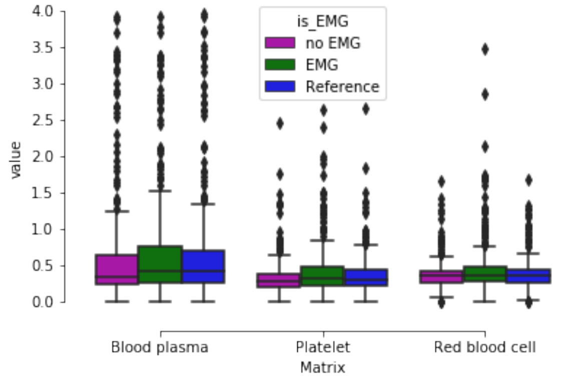

is_EMG
no EMG
EMG
Reference
value
4.0
3.5
3.0
2.5
2.0
1.5
1.0
0.5
0.0
Blood plasma
Platelet
Red blood cell
Matrix

### Table S1

Transition lists for Lipidomics, PL, v1 and v2

### Table S2

The comparison of manual and automated calibration curve fitting.

### Table S3

The description of analysed datasets. The .mzML files and the corresponding metadata are deposited in a MetaboLights[23] repository MTBLS1786.

| Identifier | Description | Method | Compounds | Number of analysed samples |
|---|---|---|---|---|
| BloodProject | Platelet, plasma and red blood cell samples derived from human whole blood (unpublished) | LC-MS/MS[24] | Polar, intracellular, extracellular | 2702 |
| ALEsKOs | Unevolved and evolved gene knockout *E. coli* populations[3] | LC-MS/MS[24] | Polar, intracellular | 1465 |
| ALEWt | End-point *E. coli* populations[25] | LC-MS/MS[24] | Polar, intracellular | 183 |
| Fermentors | Sampled from fermentors during batch and/or continuous culture at various dilution rates and various biomass concentrations (from ~0.5 to ~45 g/L) (unpublished) | LC-MS/MS[24] | Polar, intracellular | 281 |
| RapidRIP | *E. coli* samples analysed with RapidRIP method[18] | LC-MS/MS[18] | Polar, intracellular | 278 |
| Mouse | Mouse samples (blood plasma, the peritoneum and the lung)[26] | LC-MS/MS[24] | Polar, extracellular | 253 |
| Flux | *E. coli* fluxomics samples[18,27] | LC-MS/MS[28] | Polar, intracellular | 53 |
| Lipidomics | Canine blood plasma lipidomics samples[19] | LC-MS/MS[19] | Non-polar and polar, extracellular | 117 |
